# Inhibition of OCT4 Binding at the *MYCN* Locus Induces Neuroblastoma Cell Death Accompanied by Downregulation of Transcripts with High-Open Reading Frame Dominance

**DOI:** 10.1101/2023.06.08.544289

**Authors:** Kazuma Nakatani, Hiroyuki Kogashi, Takanori Miyamoto, Taiki Setoguchi, Tetsushi Sakuma, Kazuto Kugou, Yoshinori Hasegawa, Takashi Yamamoto, Yoshitaka Hippo, Yusuke Suenaga

## Abstract

Amplification of *MYCN* is observed in high-risk neuroblastomas (NBs) and is associated with a poor prognosis. *MYCN* expression is directly regulated by multiple transcription factors, including OCT4, MYCN, CTCF, and p53 in NB. Our previous study showed that inhibition of p53 binding at the *MYCN* locus induces NB cell death. However, it remains unclear whether other transcription factors contribute to NB cell survival. In this study, we revealed that the inhibition of OCT4 binding at the *MYCN* locus, a critical site for the human-specific OCT4–MYCN positive feedback loop, induces caspase-2-mediated cell death in *MYCN*-amplified NB. We used the CRISPR/deactivated Cas9 (dCas9) technology to specifically inhibit transcription factors from binding to the *MYCN* locus in the *MYCN*-amplified NB cell lines CHP134 and IMR32. In both cell lines, the inhibition of OCT4 binding at the *MYCN* locus reduced MYCN activity. Differentially downregulated transcripts were associated with high-open reading frame (ORF) dominance score, which is associated with the translation efficiency of transcripts. These transcripts were enriched in splicing factors, including MYCN-target genes such as *HNRNPA1* and *PTBP1*. Furthermore, transcripts with high-ORF dominance were significantly associated with genes whose high expression is associated with a poor prognosis of NB. In conclusion, the inhibition of OCT4 binding at the *MYCN* locus resulted in reduced MYCN activity, which in turn led to the downregulation of high-ORF dominance transcripts and subsequently induced caspase-2-mediated cell death in *MYCN*-amplified NB cells. Therefore, disruption of the human-specific OCT4–MYCN positive feedback loop may serve as an effective therapeutic strategy for *MYCN*-amplified NB.

**Contribution to the field:** Neuroblastoma (NB) is a childhood tumor. Amplification of *MYCN* is frequently observed in high-risk NBs and is linked to a poor prognosis. Multiple transcription factors, including OCT4, MYCN, CTCF, and p53, regulate *MYCN* expression by binding to the *MYCN* locus. This study investigated the contribution of these transcription factors in NB cell survival. We used CRISPR/deactivated Cas9 (dCas9) technology to specifically inhibit transcription factors from binding to the *MYCN* locus in *MYCN*-amplified NB cell lines. We found that the inhibition of OCT4 binding at the *MYCN* locus, a critical site for the human-specific OCT4–MYCN positive feedback loop, reduces MYCN activity and induces NB cell death. A detailed investigation of the molecular mechanisms of cell death revealed that the downregulated transcripts after suppressed MYCN activity were associated with high-open reading frame (ORF) dominance scores, which are associated with translation efficiency of transcripts. These transcripts were enriched in splicing factors, including MYCN-target genes such as *HNRNPA1* and *PTBP1*. Reduced expression of these splicing factors altered the *PKM* mRNA splicing accompanied by the induction of p53–caspase-2–MDM2-mediated cell death. These findings suggest that disrupting the human-specific OCT4–MYCN positive feedback loop may serve as a promising therapeutic strategy for *MYCN*-amplified NB.

## 1 Introduction

Neuroblastoma (NB) is the most common extracranial solid tumor in children, accounting for 12%– 15% of all cancer-related deaths in children (1–3). At least 40% of all NBs are designated as high-risk tumors and often show *MYCN* amplification (4). Amplification of *MYCN* is observed in 25% of high-risk cases and correlates with poor clinical outcomes in patients with NB (5,6). *Th-MYCN* mice, which are used as a preclinical *in vivo* model of NB, spontaneously develop NB, highlighting the significance of *MYCN* as a potent oncogene in the pathogenesis of NB (7). Despite current therapeutic advances, therapeutic strategy for targeting MYCN remains a medical challenge (8). Therefore, new MYCN-targeting therapeutic strategies are required to further improve patient outcomes.

MYCN, a basic helix–loop–helix transcription factor, directly regulates the transcription of genes involved in diverse cellular processes, such as cell growth, apoptosis, and differentiation (4). It directly binds to its own intron 1 region and upregulates its own expression and its *cis*-antisense gene *NCYM* by forming a positive autoregulatory loop in NB cells (9–11). In addition to MYCN, other transcription factors bind to the *MYCN* locus to regulate *MYCN* expression in NB. For example, OCT4, a transcription factor that maintains cancer stemness, is highly expressed in NB, regulates multipotency, and contributes to drug-resistant phenotypes of NB (12–17). In our previous study, we found that OCT4 stimulates *MYCN* transcription by binding to the intron 1 of *MYCN* locus, whereas MYCN stimulates *OCT4* transcription by binding to the *OCT4* promoter region (17). The OCT4-binding sequence in intron 1 of *MYCN* is not present in mice but mostly conserved in other mammals (17). In contrast, the E-box in the MYCN-binding region of the *OCT4* promoter is specific to humans and absent even in chimpanzees (17). Thus, OCT4 and MYCN form a human-specific positive feedback loop in NB (17). This human-specific positive feedback loop contributes to the stemness of *MYCN*-amplified NB by maintaining the expression of stem cell-related genes including *LIN28, NANOG*, and *SOX2* (17). Additionally, CCCTC-binding factor (CTCF), an insulator protein that is capable of regulating gene expression, stimulates *MYCN* transcription by binding to the *MYCN* promoter region (18). Moreover, we previously reported that the tumor-suppressive transcription factor p53 binds to *MYCN* exon 1 region and weakly represses *MYCN* and *NCYM* transcription (19). Blocking the p53-binding site using the CRISPR/deactivated cas9 (dCas9) system upregulates *MYCN, NCYM*, and p53 expression, inducing apoptotic cell death accompanied by caspase-2 activation (19). Thus, the p53-mediated repression of *MYCN*/*NCYM* contributes to the survival of *MYCN*-amplified NB cells (11,19). However, it remains unclear whether the regulation of *MYCN* expression by other transcription factors (OCT4, MYCN, and CTCF) contributes to NB cell survival.

In this study, we evaluated the significance of transcription factors that bind to the *MYCN* locus in NB cells. Our results suggest that the human-specific MYCN–OCT4 positive feedback loop plays a crucial role in *MYCN*-amplified NB cell survival.

## 2 Material and Methods

### 2.1 Cell culture

Human NB cell lines CHP134 and IMR32 were maintained in RPMI-1640 (Nacalai Tesque, Kyoto, Japan) supplemented with 10% fetal bovine serum (Thermo Fisher Scientific, Waltham, MA), 50 U/mL penicillin, and 50 μg/mL streptomycin (Thermo Fisher Scientific, Waltham, MA).

Neuroblastoma cell line SK-N-AS was maintained in Dulbecco’s Modified Eagle Medium (DMEM) (Sigma-Aldrich, St. Louis, MO) supplemented with 10% fetal bovine serum (Thermo Fisher Scientific, Waltham, MA), 50 U/mL penicillin, and 50 μg/mL streptomycin (Thermo Fisher Scientific, Waltham, MA).

### 2.2 Vector construction

To inhibit transcription factor binding at the *MYCN* locus, we designed CRISPR guide RNAs against the MYCN-binding site (9,10), OCT4-binding site (17), CTCF-binding site A (18), p53-binding site (19), and CTCF-binding site B (data from the UCSC Genome Browser). A CRISPR/dCas9 vector was constructed as follows: pX330A_dCas9-1×2 (Addgene, Watertown, MA; plasmid ID 63596) (20) was treated with BpiI (Thermo Fisher Scientific, Waltham, MA). Thereafter, annealed oligonucleotides (p53-binding site: sense: 5′-CACCGCGCCTGGCTAGCGCTTGCT-3′, antisense: 5′-AAACAGCAAG CGCTAGCCAGGCGC-3′; OCT4-binding site: sense: 5′-CACC AGCAGGGCTTGCAAACCGCC-3′, antisense: 5′-AAACGGCGGTTTGCAAGCCCTGCT-3′; MYCN-binding site: sense: 5′-CACC GGGAGGGGGCATGCAGATGC-3′, antisense: 5′-AAAC GCATCTGCATGCCCCCTCCC-3′; CTCF-A-binding site: sense: 5′-CACC TCTCCGCGAGGTGTCGCCTT-3′, antisense: 5′-AAACAAGGCGACACCTCGCGGAGA-3′; and CTCF-B-binding site: sense: 5′-CACCCCAGCAGGCGGCGATATGCG-3′, antisense: 5′-AAACCGCATATCGCCGCCTGCTGG-3′) were inserted into the digested vector.

### 2.3 Transfection

Plasmid transfection was performed using the Neon Transfection System (Invitrogen, Carlsbad, CA) according to the manufacturer’s instructions. We used 2 × 10^5^ cells and 4 μg of the plasmid per transfection. When performing the CUT&RUN assay and RNA isolation for quantitative real-time reverse transcription-polymerase chain reaction (qRT-PCR), plasmid transfections were performed using Lipofectamine 3000 transfection reagent (Invitrogen, Carlsbad, CA), according to the manufacturer’s instructions.

### 2.4 WST assay

Cell viability was evaluated using the Cell Counting Kit-8 (CCK-8; Dojindo Laboratories, Kumamoto, Japan), according to the manufacturer’s protocol. Briefly, 100 μL of dCas9-transfected cell suspension (5,000 cells/well) was seeded in a 96-well plate. Ninety-six hours after transfection of CRISPR/dCas9, 10 μL of CCK-8 reagent was added into each well of the 96-well plate, and then, the cells were incubated for 2 h at 37°C in a 5% CO_2_ incubator. Cell proliferation was monitored at 450 nm using CORONA absorbance microplate reader (MTP-310, CORONA ELECTRIC, Ibaraki, Japan).

### 2.5 Cytotoxicity assay

To evaluate cell damage, we measured lactate dehydrogenase (LDH) activity released from cells. LDH activity was measured using the LDH Cytotoxicity Assay Kit (Nacalai Tesque, Kyoto, Japan), according to the manufacturer’s instructions. Briefly, 100 μL of dCas9-transfected cell suspension (10,000 cells/well) was seeded in a 96-well plate. Ninety-six hours after transfection of CRISPR/dCas9, 100 μL of the substrate solution was added into each well of the 96-well plate. After which, the cells were incubated for 20 min at room temperature under shading condition, and then, 50 μL of the stop solution was added into each well of the 96-well plate. LDH activity was monitored at 490 nm using 2030 ARVO X (PerkinElmer, Kanagawa, Japan).

### 2.6 CUT&RUN assay

Twenty-four hours after the transfection of CRISPR/dCas9, CUT&RUN (CUT&RUN Assay Kit, #86652, Cell Signaling Technology, Danvers, MA) was performed according to the manufacturer’s instructions. The following antibodies were used in the assay: anti-OCT4 antibody (15 μL/assay, #2750; Cell Signaling Technology, Danvers, MA) and Rabbit (DA1E) mAb IgG XP^®^ Isotype Control (15 μL/assay, #66362; Cell Signaling Technology, Danvers, MA). DNA obtained from the CUT&RUN assay was amplified using SYBR Green qRT-PCR with the StepOnePlus™ Real-Time PCR System (Thermo Fisher Scientific, Waltham, MA). The following primer set was used: forward 5′-TCCTGGGAACTGTGTTGGAG-3′ and reverse 5′-CTCGGATGGCTACAGTCTGT -3′. The detected DNA levels were normalized by the input signal.

### 2.7 RNA isolation and qRT-PCR

One day after CRISPR/dCas9 transfection, the total RNA from dCas9-transfected NB cells was isolated using the RNeasy Mini Kit (Qiagen, Hilden, Germany) following the manufacturer’s instructions. cDNA was synthesized using SuperScript II with random primers (Invitrogen, Carlsbad, CA). qRT-PCR was performed using SYBR Green PCR with the StepOnePlus™ Real-Time PCR System (Thermo Fisher Scientific, Waltham, MA). The following primer sets were used: *MYCN*, forward: 5′-TCCATGACAGCGCTAAACGTT-3′ and reverse: 5′-GGAACACACAAGGTGACTTCAACA-3′ and *GAPDH*, forward: 5′-GTCTCCTCTGACTTCAACAGCG-3′ and reverse: 5′-ACCACCCTGTTGCTGTAGCCAA-3′. *β-Actin* expression was quantified using the TaqMan real-time PCR assay. The mRNA level of *MYCN* was normalized by *β-Actin* and *GAPDH*.

### 2.8 Long-read and short-read RNA-sequencing

Twenty-four hours after CRISPR/dCas9 transfection, the total RNA from dCas9-transfected NB cells was isolated using the RNeasy Mini Kit (Qiagen, Hilden, Germany) following the manufacturer’s instructions. An Iso-Seq library was prepared as described in the Procedure & Checklist-Iso-Seq Express Template Preparation for Sequel and Sequel II Systems, Version 02, October 2019 (Pacific Biosciences, Menlo Park, CA). Briefly, cDNA was synthesized and amplified using the NEBNext Single Cell/Low Input cDNA Synthesis & Amplification Module (New England Biolabs, Ipswich, MA), Iso-Seq Express Oligo Kit (Pacific Biosciences, Menlo Park, CA), and barcoded primers. The size of the amplified cDNA was selected using ProNex beads (Promega, Madison, WI) under standard conditions. The Iso-Seq library was prepared from the size-selected cDNA using SMRTbell Express Template Prep Kit 2.0 (Pacific Biosciences, Menlo Park, CA). The Iso-Seq libraries were sequenced on the PacBio Sequel IIe with Sequel ICS v11.0 for 24 h using a single cell of Sequel II SMRT Cell 8M Tray, Sequel II Sequencing Kit 2.0, Sequel II Binding Kit 2.1, and Internal Control 1.0 (Pacific Biosciences, Menlo Park, CA). Circular consensus sequencing (CCS) reads were created using this instrument. An RNA-sequencing (RNA-seq) library was prepared using the NEBNext rRNA Depletion Kit v2 (Human/Mouse/Rat) and the NEBNext Ultra II Directional RNA Library Prep Kit for Illumina (New England Biolabs, Ipswich, MA). The RNA-Seq libraries were sequenced on NextSeq 500 using the NextSeq 500/550 High Output Kit v2.5 (75 cycles) (Illumina, San Diego, CA).

### 2.9 Bioinformatic analysis

Demultiplexing of CCS reads and removal of cDNA primers were performed using the lima command of SMRT Tools v11.0 (Pacific Biosciences, Menlo Park, CA) with the parameters of Iso-Seq data. The polyA tail and artificial concatemer reads were removed using the isoseq3 refine.

High-quality isoforms were obtained using the isoseq3 cluster with the parameter --use-qvs. To collapse the transcripts using the isoseq3 collapse command, the high-quality isoform reads were aligned to the human genome GRCh38 using minimap2 v2.24 (21) with the parameter --preset ISOSEQ. To remove 3’-end intrapriming artifact and RT-switching artifact, quality control and filtering were performed using SQANTI3 (22) with the genome annotation (Ensembl GRCh38 release105). Novel isoforms were identified from the filtered transcripts of all samples using the TALON v5.0 pipeline (23) with the parameter --cov 0.95 --identity 0.95 --observed. Transcript reference sequences, including novel and known transcript sequences, were created using SQANTI3 and used in the following short-read RNA-seq analysis. Salmon v1.9.0 was used to quantify transcript expression levels with the parameter fldMean 260 --fldSD 73. Differentially expressed transcripts were analyzed using the high-throughput gene expression data analysis tool DIANE (https://diane.bpmp.inrae.fr/) (24). Differentially expressed transcripts were filtered by setting the log_2_ fold change (sgRNA OCT4/no sgRNA) to 0.58 and false discovery rate (FDR) to 0.05 as threshold values.

### 2.10 Functional annotation analysis

DAVID (https://www.david.ncifcrf.gov) (25) was used to identify the enriched molecular functions and pathways related to the genes of interest. *Q*-values (*P*-values adjusted for FDR) were calculated using the Benjamini–Hochberg method in DAVID.

Enrichr (http://amp.pharm.mssm.edu/Enrichr/) (26–28) was used to analyze the enriched molecular functions and pathways related to the differentially downregulated genes after OCT4-binding inhibition. “ENCODE and ChEA Consensus TFs from ChIP-X”, “TF Perturbations Followed by Expression”, and “ENCODE TF ChIP-seq 2015” were used as gene-set libraries. *Q*-values (*P*-values adjusted for FDR) were calculated using the Benjamini–Hochberg method in Enrichr.

### 2.11 Kaplan–Meier analysis-based prognosis classification of transcripts

Genes detected in the long-read RNA-seq analysis of CHP134 and IMR32 (9,144 genes) were input into R2 Genomics Analysis and Visualization Platform (http://r2.amc.nl, Tumor Neuroblastoma -Kocak -649 -custom -ag44kcwolf, GSE45547) for Kaplan–Meier analysis to extract a set of genes associated with a poor prognosis of NB. For the type of survival, we selected overall survival. *Q*-values (*P*-values adjusted for FDR) were calculated using the Benjamini–Hochberg method in R2.

### 2.12 Western blot analysis

The cells were lysed with RIPA buffer (50 mmol/L Tris-HCl buffer (pH 7.6), 150 mmol/L NaCl, 1(w/v)% Nonidet P40 Substitute, 0.5(w/v)% sodium deoxycholate, protease inhibitor cocktail, and 0.1(w/v)% SDS; # 08714-04, Nacalai Tesque, Kyoto, Japan) and benzonase (Merck Millipore, Billerica, MA) and MgCl_2_ at final concentrations of 25 U/μL and 2 mM, respectively; incubated at 37°C for 1 h; and centrifuged at 10,000 × *g* for 10 min at 4°C. Thereafter, the supernatant was collected and denatured in SDS sample buffer (125 mM Tris-HCl, pH 6.8, 4% SDS, 10% sucrose, 0.01% BPB, and 10% 2-mercaptoethanol). Cellular proteins were resolved using sodium dodecyl sulfate-polyacrylamide gel electrophoresis before being electroblotted onto polyvinylidene fluoride membranes (#1704156, Bio-Rad Laboratories, Hercules, CA). The membranes were incubated with the following primary antibodies for 60 min at room temperature: anti-Cas9 (1:1000 dilution; #14697S, Cell Signaling Technology, Danvers, MA), anti-MDM2 (1:1000 dilution; OP46, Merck Millipore, Billerica, MA), anti-p53 (1:1000 dilution; #2524, Cell Signaling Technology, Danvers, MA), anti-caspase-2 (1:1000 dilution; sc-5292, Santa Cruz Biotechnology, Dallas, TX), anti-caspase 3 (1:1000 dilution; sc-7148, Santa Cruz Biotechnology, Dallas, TX), and anti-actin (1:1000 dilution; FUJIFILM Wako Pure Chemical Corporation, Osaka, Japan). The membranes were then incubated with horseradish peroxidase-conjugated secondary antibodies (anti-rabbit IgG at 1:5000 dilution or anti-mouse IgG at 1:5000 dilution; both from Cell Signaling Technology, Danvers, MA), and the bound proteins were visualized using a chemiluminescence-based detection kit (ImmunoStar Zeta; ImmunoStar LD, FUJIFILM Wako Pure Chemical Corporation, Osaka, Japan). Chemiluminescence was detected using ImageQuant™ LAS4000 (GE Healthcare, Chicago, IL).

To detect MYCN protein expression, western blotting was performed using an Abby instrument (ProteinSimple, Tokyo, Japan) with 25-min separation at 375 V, 10-min blocking, 30-min primary antibody incubation (anti-MYCN antibody, 1:100 dilution; #9405, Cell Signaling Technology, Danvers, MA), and 30-min secondary antibody incubation (DM-001, ProteinSimple, Tokyo, Japan). RePlex™ Module (RP-001, ProteinSimple, Inc., Tokyo, Japan) was used to detect total proteins.

Default assay parameters were used for data analysis, and peak areas were calculated using the Compass software (ProteinSimple, Tokyo, Japan). The peak area values were used to represent the intensity of target proteins. MYCN intensity was normalized by the total protein intensity.

### 2.13 Open reading frame dominance score analysis

The transcript sequences detected using long-read RNA-seq analysis were used to calculate ORF dominance, as previously described (29,30).

### 2.14 Statistical analysis

Statistical analysis software “R” was used for data analysis. Mann–Whitney *U*-test, Student’s *t*-test, and Kruskal–Wallis test were performed as appropriate. A *p*-value < 0.05 was considered statistically significant.

## 3 Results

### 3.1 CRISPR/dCas9 targeting transcription factor-binding sites at the *MYCN* locus reduced the viability in *MYCN*-amplified NB cells

Deactivated Cas9 (dCas9) disrupts the binding of transcription factors to specific sites (31). To inhibit transcription factor binding at the *MYCN* locus, we designed CRISPR guide RNAs against the MYCN-binding site (9,10), OCT4-binding site (17), CTCF-binding site A (18), p53-binding site (19), and CTCF-binding site B (data from the UCSC Genome Browser) (Figure 1A). A previous study has demonstrated that CTCF binds to the upstream region of *MYCN* and promotes its transcription (18).

**Figure 1.**
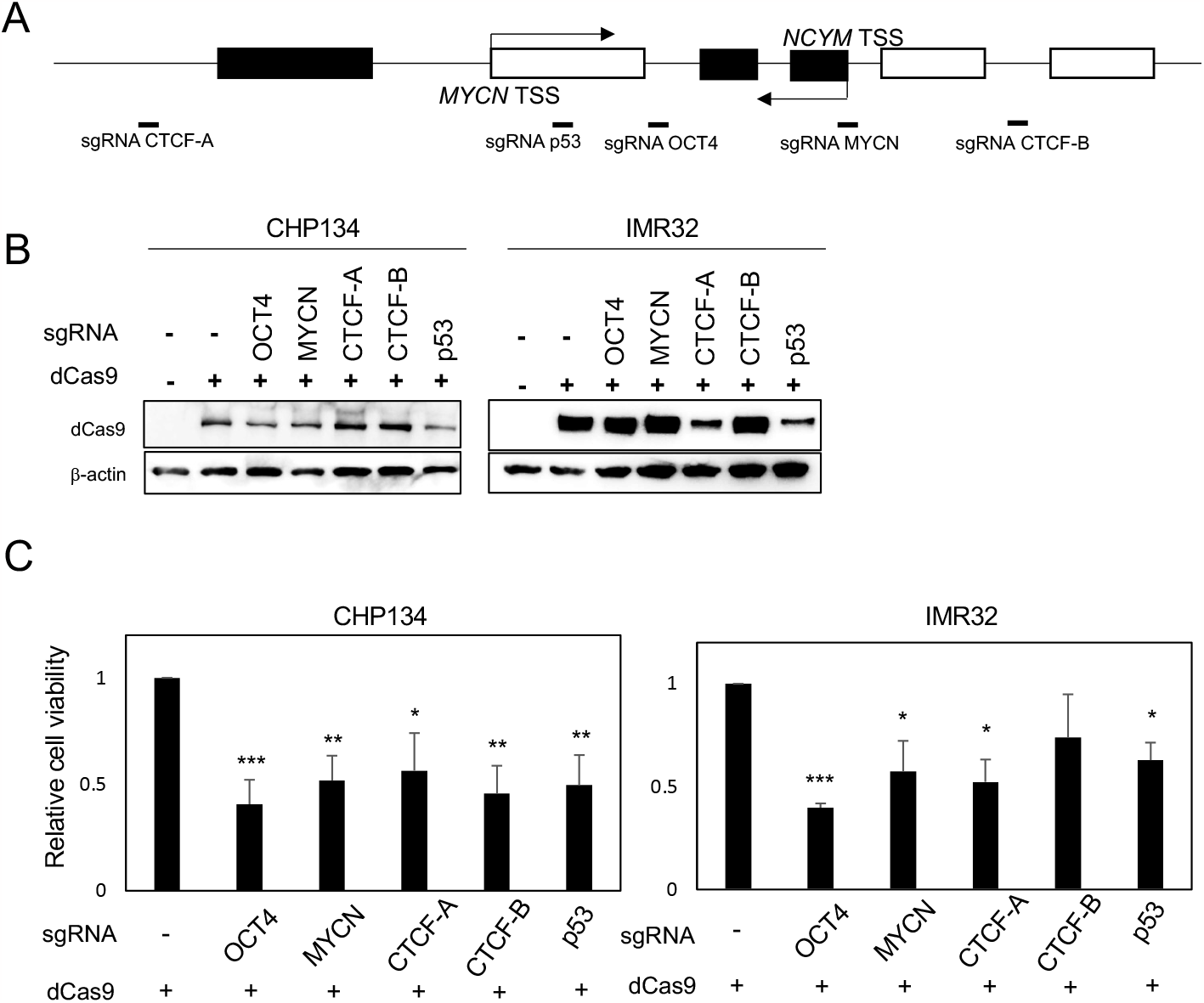
CRISPR/dCas9 targeting the *MYCN* locus reduces the viability of *MYCN*-amplified neuroblastoma cells. (A) A diagram of the *MYCN*/*NCYM* locus with the positions of targeting sgRNAs. The white and black boxes indicate the *MYCN* and *NCYM* regions, respectively. TSS: transcription start site. (B) Western blotting of dCas9 protein in CHP134 and IMR32 cells. Twenty-four hours after CRISPR/dCas9 transfection, these cells were subjected to western blotting. *β-Actin* was used as a loading control. (C) Ninety-six hours after CRISPR/dCas9 transfection, the viability of CHP134 and IMR32 was measured using the WST assay. *: *p* < 0.05; **: *p* < 0.01; ***: *p* < 0.001. Data were analyzed using student’s *t-*test (compared with no sgRNA). Error bars represent SEM of six independent experiments.

However, using the UCSC Genome Browser, we discovered an additional CTCF-binding site located within the gene body of *MYCN* (Figure S1), whose function has not been investigated in previous study (18). Therefore, we designed CRISPR guide RNAs for both CTCF-binding sites. For convenience, we designated the CTCF-binding site upstream of *MYCN* as CTCF-A and the gene body region as CTCF-B (Figures 1A and S1). We transfected all-in-one CRISPR/dCas9-sgRNA vectors into CHP134 and IMR32 cells, both of which are *MYCN*-amplified NB cells (Figure 1B).

The viability of CHP134 and IMR32 cells was significantly reduced by dCas9 targeting the OCT4-binding site, MYCN-binding site, CTCF-A site, and p53-binding site (Figure 1C). Among these targets, dCas9 targeting the OCT4-binding site most significantly decreased in cell viability in both *MYCN*-amplified NBs (CHP134 and IMR32) (Figure 1C). On the other hand, the viability of *MYCN*-nonamplified NB cells (SK-N-AS) was not affected by dCas9 targeting the OCT4-binding site (Figure S2).

### 3.2 Inhibition of OCT4 binding at the *MYCN* locus suppresses MYCN activity

As cell viability was significantly reduced by dCas9 targeting the OCT4-binding site in both *MYCN*-amplified NB cell lines (Figure 1C), we investigated the effect of inhibition of the OCT4-binding site on MYCN activity. Twenty-four hours after dCas9 transfection, CRISPR/dCas9 inhibited OCT4 binding at the *MYCN* locus (Figure 2A, B) and suppressed the expression of *MYCN* mRNA compared to control (dCas9 without sgRNA) (Figure 2C). To identify the transcriptomic changes after OCT4-binding inhibition, we performed short-read RNA-seq combined with long-read RNA-seq of CHP134 and IMR32 cells 24 h after dCas9 transfection. We have listed the detected transcripts and their expression levels in Table S1. Through this analysis, we detected 17,601 annotated transcripts (transcript ID starts with ENST∼) and 70,753 unannotated transcripts (transcript ID starts with TALONT∼) in the combined CHP134 and IMR32 cell samples. Notably, the number of unannotated transcripts was approximately four times higher than the number of annotated transcripts. For example, we detected 1 annotated transcript (Figure S3A (ii)) and 4 unannotated transcripts (Figure S3A (i), (iii), (iv), and (v)) transcribed from the *MYCN* locus, and 2 annotated transcripts (Figure S3A (viii) and (ix)) and 2 unannotated transcripts (Figure S3A (vi) and (vii)) transcribed from the *NCYM* locus using long-read RNA-seq analysis. *NCYM* non-coding RNA (ENST00000641263), previously reported to function as a non-coding RNA in NB (32), was not detected in this study. The normalized expression counts analyzed from short-read RNA-seq of *MYCN* and *NCYM* transcripts in CHP134 cells are presented in Figure S3B and C. Among these transcripts, the expression of ENST00000281043 (Figure S3A (ii)), encoding the MYCN protein, tended to be suppressed (Figure S3B), whereas that of TALONT000261009 (Figure S3A (iii)), which lacks the MYCN-coding sequence, was upregulated by OCT4-binding inhibition (Figure S3B). Western blotting showed decreased expression of the MYCN protein at 72 h after dCas9 transfection (Figure 2D). Consistent with this observation, Enrichr analysis (http://amp.pharm.mssm.edu/Enrichr/) (26–28) revealed that differentially downregulated genes were enriched in MYCN-target genes (GSE80397: downregulated

**Figure 2.**
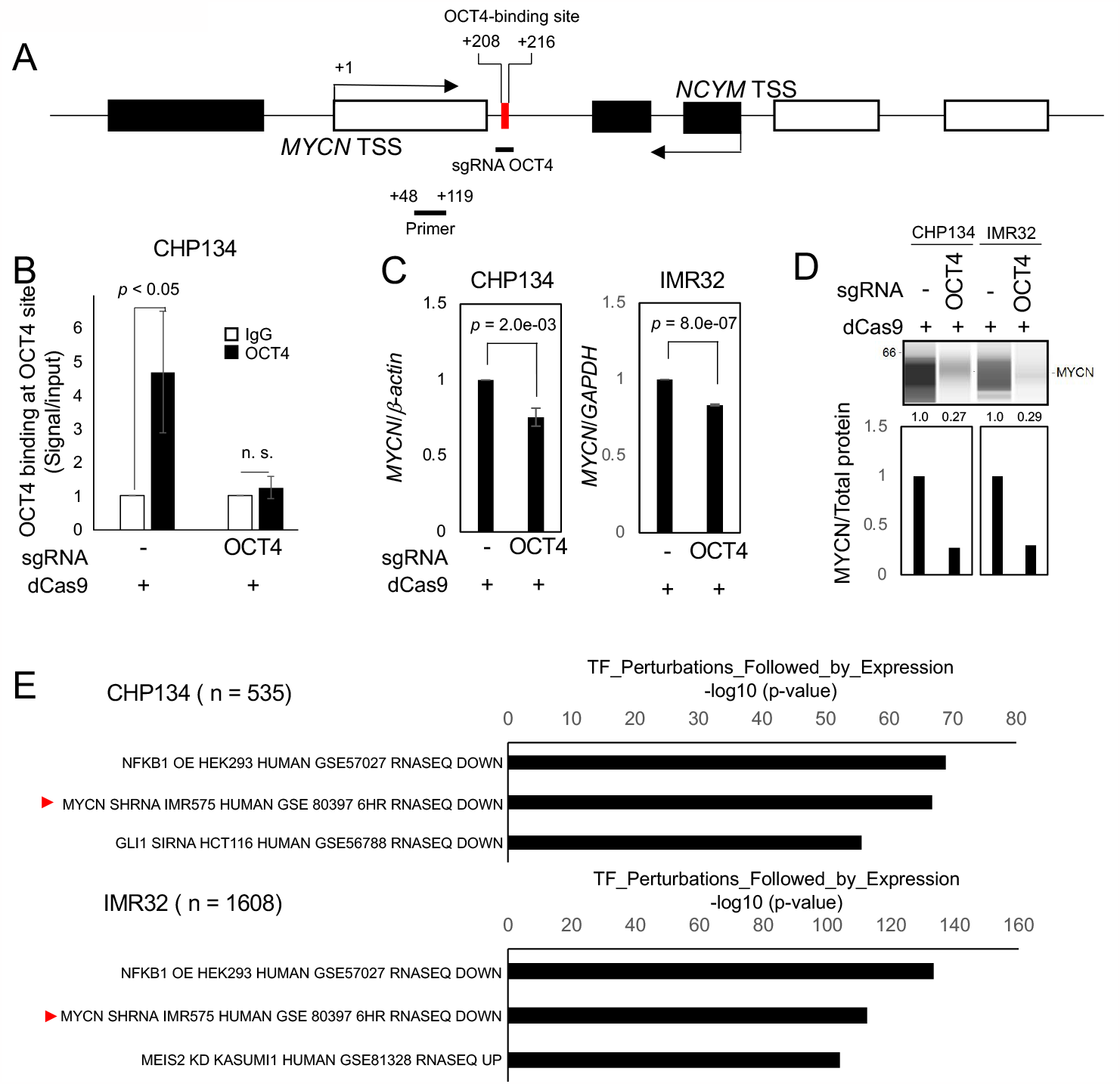
Inhibition of OCT4 binding at the *MYCN* locus reduces MYCN activity. (A) Schematic depiction of the *MYCN*/*NCYM* locus with the location of the primer used in the CUT&RUN assay. The OCT4-binding site is indicated with a red box. The white and black boxes indicate the *MYCN* and *NCYM* regions, respectively. (B) OCT4 binding at the *MYCN* locus was inhibited using CRISPR/dCas9. Twenty-four hours after the transfection of CRISPR/dCas9 targeting the OCT4-binding site, CHP134 cells were subjected to the CUT&RUN assay. Genomic DNA was amplified using quantitative real-time reverse transcription-polymerase chain reaction (qRT-PCR) using primer in Figure 2A. The signals were normalized by input signals. IgG was used as an isotype control. Error bars represent SEM of four independent experiments. Data were analyzed using student’s *t*-test. (C) qRT-PCR analyses of *MYCN* in CRISPR/dCas9-transfected CHP134 and IMR32 cells. One day after transfection, mRNA expression levels were measured using qRT-PCR with *β-actin* or *GAPDH* as an internal control. Data were analyzed using Student’s *t-*test. Error bars represent SEM of three independent experiments. (D) Quantitative analysis of western blot was performed on CHP134 and IMR32 cells transfected with CRISPR/dCas9. MYCN protein expression levels were measured using the SimpleWestern™ system at 72 h after transfection. The upper panel displays the MYCN protein band processed by the SimpleWestern™ system, whereas the lower panel shows the quantified MYCN signal normalized by total protein level. Data is representative data. (E) Differentially downregulated genes after inhibition of OCT4 binding at the *MYCN* locus were enriched in MYCN*-*target genes. Enrichr analysis (http://amp.pharm.mssm.edu/Enrichr/) summary of enriched transcription factor-target genes.

### 3.3 Disruption of OCT4–MYCN Axis Induces Neuroblastoma Cell Death

gene set after *MYCN* knockdown in IMR575) (Figure 2E) and MYC/MAX-target genes (Table S2). On the contrary, in the Enrichr analysis of three independent gene-set libraries (ENCODE and ChEA Consensus TFs from ChIP-X, TF Perturbations Followed by Expression, and ENCODE TF ChIP-seq 2015), enrichment of OCT4-target genes was not observed, suggesting no off-target effects of CRISPR/dCas9 on the expression of other OCT4-target genes (Table S2). These findings indicate that CRISPR/dCas9 specifically inhibited OCT4 binding at the *MYCN* locus and suppressed MYCN activity in *MYCN*-amplified NB.

### 3.4 Inhibiting OCT4 binding at the *MYCN* locus downregulates high-ORF dominance transcripts associated with poor prognosis

We examined how the reduced MYCN activity altered the NB transcriptome. In our previous study, we developed the ORF dominance score, which is defined as the fraction of the longest ORF in the sum of all putative ORF lengths within a transcript sequence (29). This score correlates with translation efficiency of coding transcripts and non-coding RNAs (29). Our previous *in silico*-based analysis suggested that noncoding transcripts with high-ORF dominance are associated with downstream genes of MYCN in humans (29). Therefore, we investigated whether MYCN functions as a regulator of transcripts with high-ORF dominance in NB. We calculated ORF dominance scores of differentially downregulated transcripts using long-read RNA-seq analysis (Table S3). The differentially downregulated transcripts had significantly higher ORF dominance than all transcripts, and this trend was observed for both coding and non-coding RNAs (Figure 3A). Additionally, isoform expression analysis from short-read RNA-seq showed similar results, revealing that the differentially downregulated transcripts had significantly higher ORF dominance in both coding and non-coding transcripts (Figure S4). These findings indicate that MYCN maintains the expression of transcripts with high-ORF dominance in NB.

**Figure 3.**
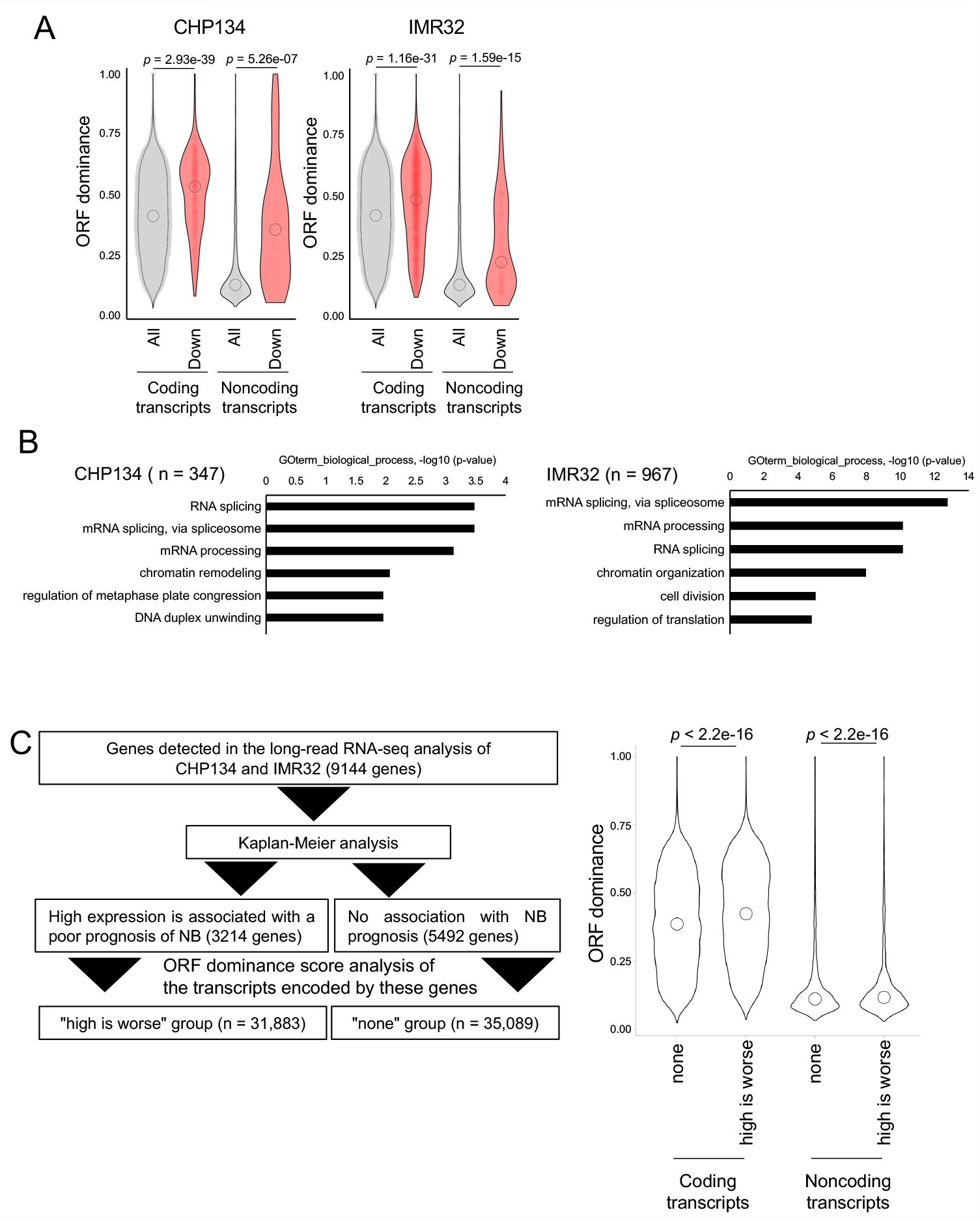
Downregulated transcripts after dCas9 transfection are associated with high-open reading frame (ORF) dominance score in neuroblastoma cells. (A) Differentially downregulated transcripts were associated with high-ORF dominance in CHP134 (left) and IMR32 cells (right). The number of samples was as follows: coding transcripts (CHP134: all, n = 51,400; down, n = 610, IMR32: all, n = 53,245; down, n = 2,047). Noncoding transcripts (CHP134: all, n = 3,464; down, n = 29, IMR32: all, n = 3,601; down, n = 144). The summary of the data is shown as a violin plot reflecting the data distribution and an open circle indicating the median of the data. *P*-values were calculated using Mann–Whitney *U*-test. (B) Gene Ontology (GO) analysis of differentially downregulated transcripts with high-ORF dominance (ORF dominance > 0.5). (C) The “high is worse” transcripts (coding, n = 30,470; noncoding, n = 1,413) showed higher ORF dominance than the “none” transcripts (coding, n = 31,895; noncoding, n = 3,194). The “high is worse” group contains transcripts whose high expression is associated with a poor prognosis of neuroblastoma. The “none” group transcripts are not associated with the prognosis of neuroblastoma. A summary of the data is shown as a violin plot reflecting data distribution and an open circle indicating the median of the data. *P*-values were calculated using Mann–Whitney *U*-test.

Next, to analyze the functions of transcripts with high ORF dominance, we extracted transcripts with high-ORF dominance (ORF dominance > 0.5) from the differentially downregulated transcripts and performed Gene Ontology (GO) analysis using DAVID. Transcripts with high-ORF dominance were associated with the GO terms “mRNA processing,” “mRNA splicing via spliceosome,” and “RNA splicing” (Figure 3B and Table S4). Notably, the genes encoding the splicing factors *HNRNPA1* and *PTBP1* are the targets of MYCN, and a decrease in MYCN activity induces the downregulation of *HNRNPA1* and *PTBP1* expression and suppresses the proliferation of *MYCN*-amplified NB cells (33). Consistent with this previous report, the expression of *HNRNPA1* and *PTBP1*, the target genes of MYCN, was downregulated after OCT4-binding inhibition in this study (Figure S5A). HNRNPA1 and PTBP1 regulate the alternative splicing of the pyruvate kinase gene (*PKM*) and facilitate the switch from the canonical isoform *PKM1* to the cancer-related isoform *PKM2* (33,34). The knockdown of *PTBP1, HNRNPA1*, and their downstream target *PKM2* represses the proliferation of *MYCN*-amplified NB (33). Similarly, the *PKM2*/*PKM1* ratio was significantly decreased by OCT4-binding inhibition in this study (Figure S5B), suggesting that the splicing switch from *PKM1* to *PKM2* underlies the mechanism of inhibition of NB proliferation after transfection of CRISPR/dCas9 targeting the OCT4-binding site.

We examined the expression of cell death-related proteins using western blotting to gain insights into the mechanism of inhibition of NB proliferation. In our previous study, we found that blocking the p53-binding site at the *MYCN* locus using CRISPR/dCas9 results in the cleavage of caspase-2 and MDM2 and induction of p53 expression (19), which is associated with the p53–MDM2–caspase-2 positive feedback loop (35). Consistent with this report, cleavage of caspase-2 and MDM2, but not caspase-3, and induction of p53 expression were observed in *MYCN*-amplified NB cells at 72 h after transfection of CRISPR/dCas9 (Figure S5C). To evaluate cytotoxicity, the activity of LDH released from cells was measured using a cytotoxicity LDH assay at 96 h after transfection of CRISPR/dCas9. The LDH activity was enhanced by CRISPR/dCas9 targeting the OCT4-binding site in CHP134 and IMR32 cells (Figure S5D). This result suggests that inhibition of OCT4 binding at the *MYCN* locus induces apoptosis in NB cells via the activation of the p53–MDM2–caspase-2 positive feedback loop.

Finally, we investigated whether genes encoding transcripts with high-ORF dominance are associated with the poor prognosis of NB using the R2 Genomics Analysis and Visualization Platform. Genes detected in the long-read RNA-seq analysis of CHP134 and IMR32 (9,144 genes) were input into the Kaplan-Meier analysis of the R2 database to extract a set of genes associated with poor NB prognosis of NB. We found 3,214 genes whose high expression was associated with a poor prognosis of NB. The transcripts encoded by these genes were classified into “high is worse” group (n = 31,883) (Table S5). We also identified 5,492 genes that were not associated with NB prognosis. The transcripts encoded by these genes were classified into “none” group (n = 35,089) (Table S5). We found that the transcripts in the “high is worse” group had significantly higher ORF dominance than those in the “none” group (Figure 3C). These results indicate that MYCN regulates the expression of transcripts with high-ORF dominance, most of which are associated with poor prognosis of NB.

## 4 Discussion

In this study, we showed that the specific inhibition of OCT4 binding at the *MYCN* locus reduces MYCN activity and induces NB cell death. In our previous study, we found that high *OCT4* mRNA expression is associated with a poor prognosis of *MYCN*-amplified NBs, but not in *MYCN*-non-amplified NBs (17). Consistent with this finding, in this study, OCT4-binding inhibition in the intron 1 region of *MYCN* decreased the viability of *MYCN*-amplified NB cells (CHP134 and IMR32), but not that of *MYCN*-non-amplified NB cells (SK-N-AS), suggesting that the human-specific OCT4– MYCN mutual positive feedback loop is specifically required for the survival of *MYCN*-amplified NB. Recent reports indicate that POU family proteins and MYCN bind to human-specific *cis*-regulatory elements in cranial neural crest cells (36), suggesting that POU family proteins and MYCN have human-specific roles in neural crest development. In addition, *NCYM*, a *cis*-antisense gene of *MYCN*, encodes the homininae-specific oncoprotein and enhances metastasis of NBs (10) possibly by inhibiting apoptotic cell death (10,19,37) and regulating stemness related genes, including *OCT4, NANOG*, and *LIN28* (17). Together, the human-specific OCT4–MYCN positive feedback loop and the Homininae-specific oncoprotein NCYM form a human-specific regulatory network in NBs that contributes to malignant phenotypes. Our findings suggest that evolution of these network may contribute to cancer-specific characteristics, and a comparative analysis between humans and their close relative species may help clarify whether the human genetic background is more prone to cancer phenotypes.

In this study, inhibition of OCT4 binding at the *MYCN* locus suppressed *MYCN* and its downstream genes, including *HNRNPA1* and *PTBP1*, which are splicing factors. The reduction of *HNRNPA1* and *PTBP1* subsequently decreased splicing activity, leading to a decrease in the *PKM2*/*PKM1* ratio and activation of caspase-2. A previous study by Zhang et al. (33) reported that knockdown of PKM2 suppresses cell proliferation in *MYCN*-amplified NB cells (IMR5), but not in *MYCN*-non-amplified NB cells (SK-N-AS), suggesting a *MYCN*-amplified NB-dependent function for PKM2. Our observation that CRISPR/dCas9 targeting of the OCT4 binding site suppresses cell proliferation specifically in *MYCN*-amplified NB is therefore consistent with this report. However, the link between PKM2 and caspase-2 remains unclear. One possible explanation is that PKM2 interacts with the CDK1-cyclinB complex to facilitate cell cycle progression in gliomas, and knockdown of PKM2 decreases CDK1 kinase activity (38). Reduced CDK1 activity decreases the inhibitory phosphorylation level of the S340 residue of caspase-2, thereby leading to caspase-2 activation (39). Thus, suppressed *PKM2* expression may activate caspase-2 through the reduction of CDK1-cyclin B kinase activity.

The inhibition of OCT4 binding at the *MYCN* locus induced cell death, with increased p53 expression and cleavage of caspase-2 and MDM2. In NB, cleaved MDM2 (MDM2-p60) is generated by oridonin, an active diterpenoid derived from traditional Chinese medicine (40). The generation of MDM2-p60 stabilizes p53 expression and results in p53 accumulation for continuous activation (40). This oridonin-induced p53 activation promotes apoptosis and cell cycle arrest in NB cells (40). In addition to the caspase-mediated inhibition of MDM2, small-molecule inhibitors of MDM2, such as nutlin-3, MI-63, and RG7388, have been shown to exert anticancer activity in NB cells (41,42).

Notably, *MYCN* amplification or overexpression sensitizes NB cell lines with wild-type p53 to MDM2-p53 antagonists, including Nutlin-3 and MI-63 (41). Thus, caspase-2-mediated cleavage of MDM2 may play an essential role in cell death induced by inhibition of OCT4 binding at the *MYCN* locus.

Previously, we developed the ORF dominance score, defined as the fraction of the longest ORF in the sum of all putative ORF lengths (29). An *in silico*-based analysis suggested that noncoding transcripts with high-ORF dominance are associated with the downstream gene of MYCN in humans (29). However, whether MYCN regulates the expression of transcripts with high-ORF dominance has not yet been experimentally investigated. In this study, we investigated the effect of MYCN activity on the expression of transcripts with high-ORF dominance in *MYCN*-amplified NB cells. Our findings demonstrate that a reduction in MYCN activity led to a decrease in the expression of both coding and noncoding transcripts with high ORF dominance. Importantly, the present study identifies MYCN as the first experimentally validated regulator of ORF dominance. Moreover, we found that transcripts with high ORF dominance are associated with poor prognosis in NB patients, indicating that ORF dominance may serve as a novel prognostic marker in NB. However, it should be noted that the ORF dominance data obtained in this study were based on transcript sequences from cell lines (CHP134 and IMR32). Hence, future studies should investigate whether the ORF dominance score can serve as a prognostic marker in NB using patient-derived transcript data.

## 5 Conflict of Interest

The authors declare that this research was conducted in the absence of any commercial or financial relationships that could be construed as a potential conflict of interest.

## 6 Author Contributions

KN, HK, TM, KK, YHasegawa, TSakuma, TY, and YS performed the experiments and acquired and analyzed the data. KN, KK, YHasegawa, YHippo, and YS wrote the manuscript. KN, TSetoguchi, YHippo, and YS acquired the funds. YS designed and supervised the study. All authors contributed to manuscript preparation and approved the submitted version.

## 7 Funding

This work was partially supported by JST SPRING (Grant Number JPMJSP2109 to KN), e-ASIA Grant from the Japan Agency for Medical Research and Development (NO. 21jm0210092h0001 to YHippo), Grant-in-Aid for Scientific Research (C) JSPS Kakenhi Grant (No. 21K08610 to YS), Grant-in-Aid for Scientific Research (C) JSPS Kakenhi Grant (No. 20K09338 and No. 23K08511 to TSetoguchi), and Takeda Science Foundation (to YS) and the Innovative Medicine CHIBA Doctoral WISE Program (to KN) from Chiba University.

## 8 Acknowledgments

We thank Ryo Otaka, Kyoko Takahashi, Miho Kobatake, Taichi Yokoi, and Harumi Saida for their assistance in the study. We would like to thank Editage (www.editage.com) for English language editing.

## 10 Data Availability Statement

We have deposited the raw sequencing data in this study to the DNA Data Bank of Japan (DDBJ), under the accession number [PRJDB15933]. The deposited data can be accessed through the DDBJ website (http://www.ddbj.nig.ac.jp/).

## Notes

### Competing Interest Statement

The authors have declared no competing interest.

