## Supplementary figures and images for "Inhibition of OCT4 Binding at the *MYCN* Locus Induces Neuroblastoma Cell Death Accompanied by Downregulation of Transcripts with High-Open Reading Frame Dominance"

### Supplementary Figures

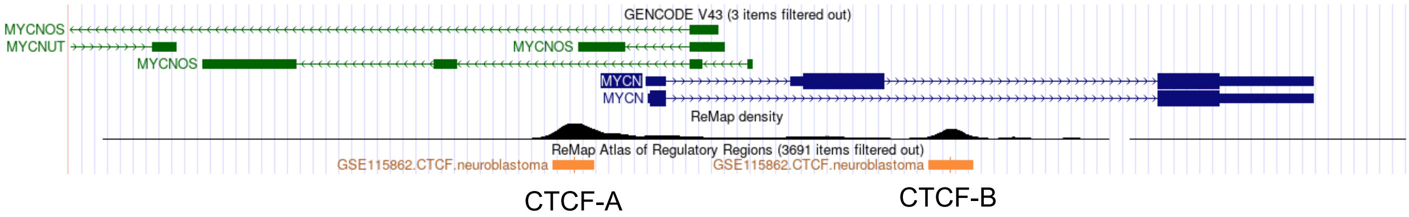

#### Figure S2

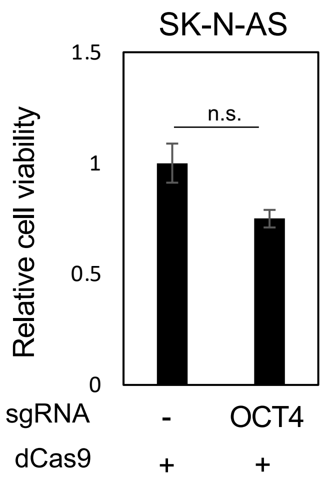

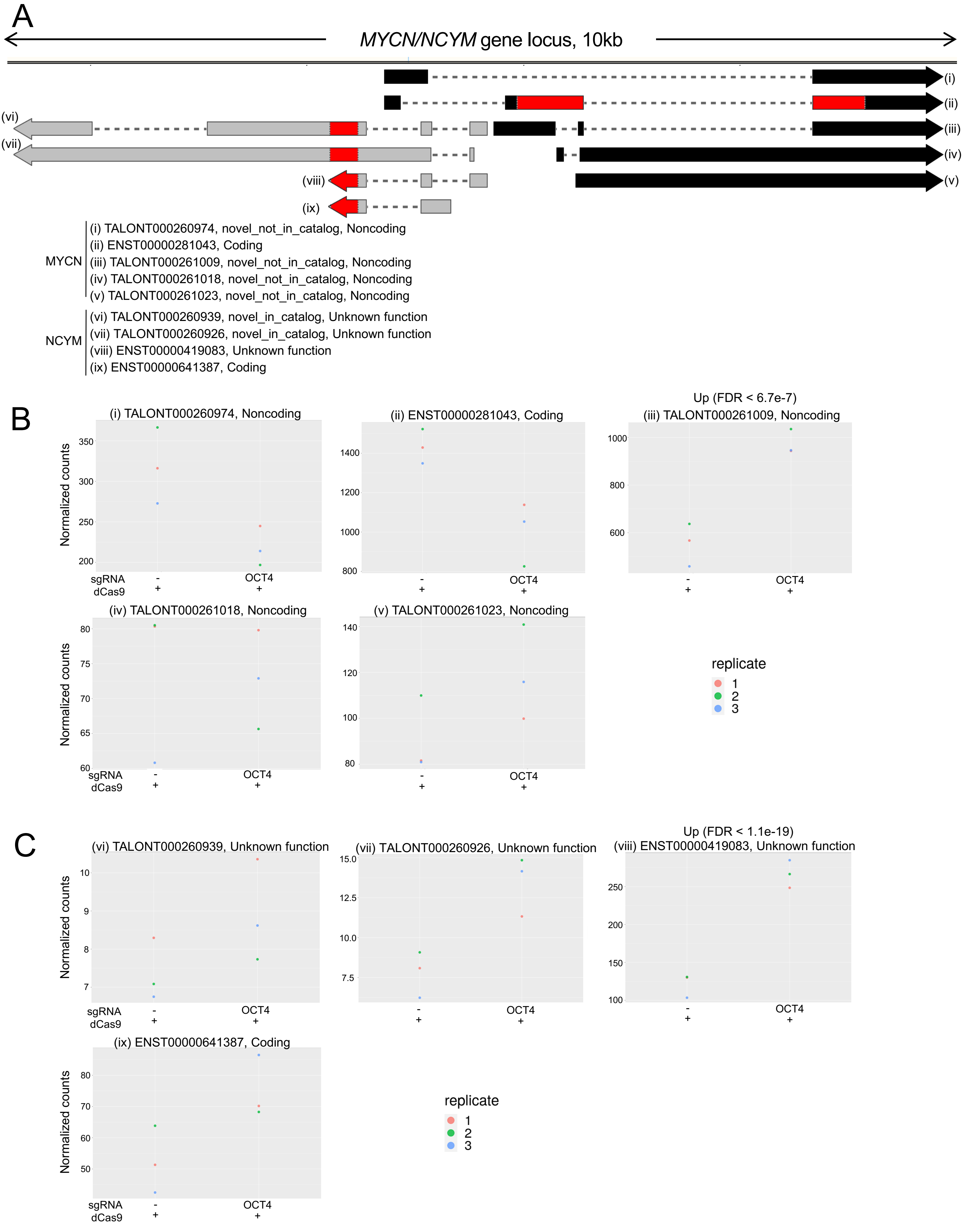

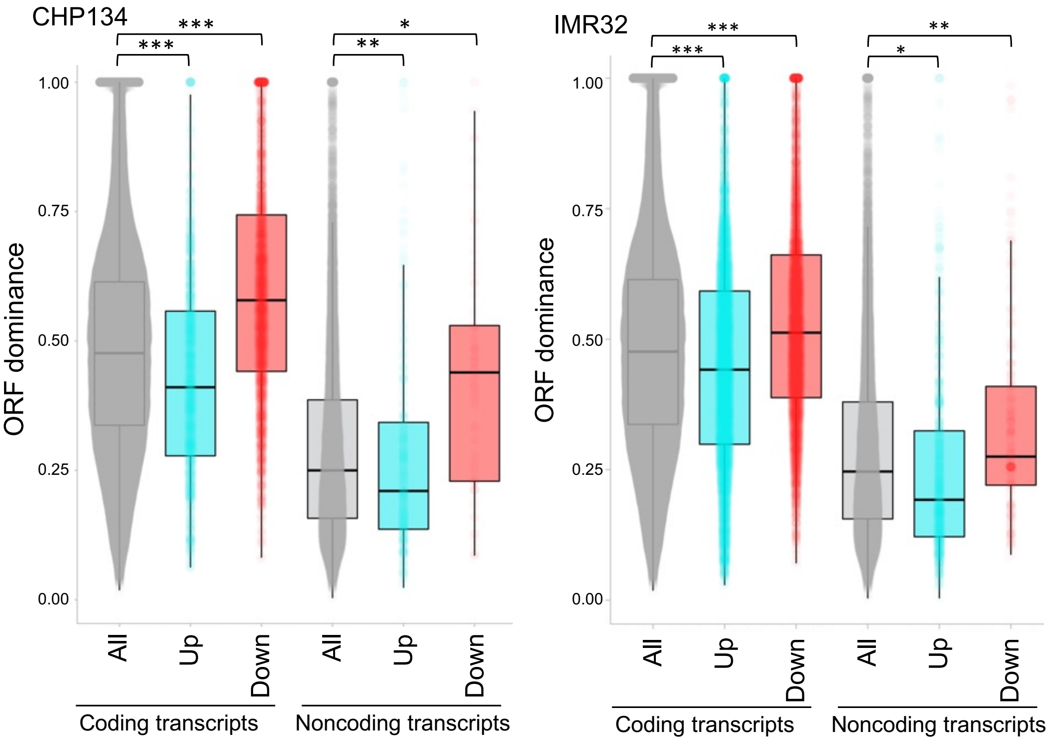

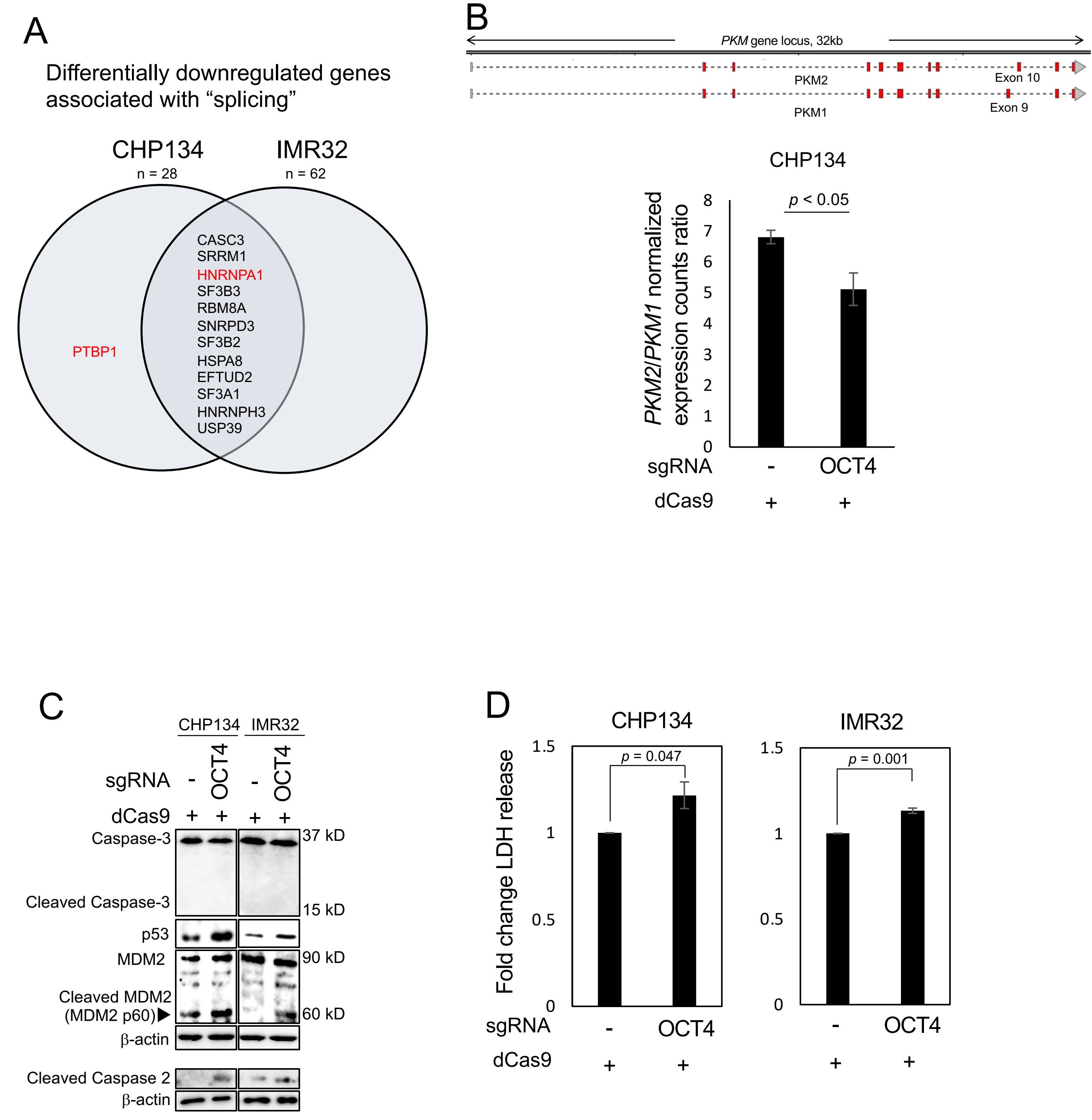
