## Supplementary Figure Legends for "Inhibition of OCT4 Binding at the *MYCN* Locus Induces Neuroblastoma Cell Death Accompanied by Downregulation of Transcripts with High-Open Reading Frame Dominance"

### 1 Supplementary Figures and Tables

**Figure S1.** CCCTC-binding factor (CTCF) enrichment at the *MYCN* locus.
UCSC Genome Browser showing the enrichment of CTCF in the upstream region of the transcription start site (TSS) (CTCF-A) and gene body (CTCF-B) of *MYCN* in GSE115862 neuroblastoma data set.

**Figure S2.** Cell viability was unchanged by deactivated cas9 (dCas9) targeting the OCT4-binding site in *MYCN* non-amplified neuroblastoma cell line (SK-N-AS).
Ninety-six hours after CRISPR/dCas9 transfection, the viability of SK-N-AS cells was measured using the WST assay. Data were analyzed using student's *t*-test (compared with no sgRNA). Error bars represent SEM (n = 3).

**Figure S3.** Transcripts detected using long-read RNA-seq analysis.
(A) A diagram of transcripts detected at the *MYCN*/*NCYM* locus. Black and gray indicate *MYCN* and *MYCNOS* (*NCYM*) transcripts, respectively. Red regions indicate coding sequences (CDS). Novel_not_in_catalog means a novel transcript not in the reference produced by a novel splice site. Novel_in_catalog means a novel transcript not in the reference produced by a known splice site. (B) Normalized expression counts (TPM) of *MYCN* transcripts from short-read RNA-seq analysis in CHP134 cells. Dots indicate biological replicates (n = 3). (C) Normalized expression counts (TPM) for *MYCNOS* (*NCYM*) transcripts from short-read RNA-seq analysis in CHP134 cells. Dots indicate biological replicates (n = 3).

**Figure S4.** Differentially downregulated transcripts had a high-open reading frame (ORF) dominance score (short-read RNA-seq analysis).
Differentially downregulated transcripts were associated with high-ORF dominance in CHP134 (left) and IMR32 (right) cells. The number of samples was as follows: coding transcripts (CHP134; all, n = 141,246, up, n =1,047, down, n = 2,286, IMR32; all, n = 142,570, up, n = 5,618, down, n = 4,255). Noncoding transcripts (CHP134; all, n = 18,328, up, n = 325, down, n = 70, IMR32; all, n = 19,691, up, n = 697, down, n = 173). A summary of data is shown as a boxplot, with the box indicating the IQR, whiskers showing the range of values that were within 1.5*IQR, and horizontal line indicating the median. *P*-values were calculated using Kruskal–Wallis test. *: $p < 1.0e-03$; **: $p < 1.0e-06$; ***: $p < 1.0e-19$

**Figure S5.** CRISPR/dCas9 targeting the OCT4-binding site induces neuroblastoma cell death accompanied by the alteration of the *PKM* mRNA splicing and activation of the p53–MDM2–caspase-2 pathway.
(A) A Venn diagram of differentially downregulated genes associated with splicing in CHP134 and IMR32 cells. (B) The *PKM2*/*PKM1* ratio was decreased by CRISPR/dCas9 targeting the OCT4-binding site. Upper panel: a diagram of transcripts detected using the long-read RNA-seq analysis at the *PKM* locus. Red regions indicate coding sequences (CDS). Lower panel: *PKM2/PKM1* normalized expression count ratio from the short-read RNA-seq analysis. (C) Western blotting of p53, MDM2, caspase-2, and caspase-3 in dCas9-transfected neuroblastoma cells. Seventy-two hours after transfection, the cells were subjected to western blotting. β-actin was used as the loading control. (D) CRISPR/dCas9 targeting the OCT4-binding site induced neuroblastoma cell death. Ninety-six hours after transfection of CRISPR/dCas9, activity of lactate dehydrogenase (LDH) released from cells was measured using the cytotoxicity LDH assay. Data were analyzed using student's *t*-test. Error bars represent SEM of three independent experiments.

**Table S1.**
The gene information including open reading frame (ORF) dominance score and normalized expression counts of each transcript (n=3 per condition).

**Table S2.**
Enrichr analysis of differentially downregulated genes after OCT4-binding inhibition in CHP134 and IMR32.

**Table S3.**
The ORF dominance of differentially downregulated transcripts.

**Table S4.**
Gene Ontology (GO) analysis of differentially downregulated transcripts with high-ORF dominance (ORF dominance > 0.5).

**Table S5.**
List of genes analyzed using the Kaplan-Meier method.
